# Identifying Integrative Molecular Pathways for Predictive Modeling of Infectious Disease

**DOI:** 10.1101/835546

**Authors:** Charles D. Schaper

## Abstract

The signs and symptoms of infectious disease are similar in presentation, such as fever and fatigue, but differ in magnitude, duration, and sequence. Although observable responses of dysfunction are well characterized, the integrative system mechanisms driving such trajectories are poorly known, even during normative circumstances. Here, molecular pathways are presented that enable predictive modeling of autonomic dysfunction due to infectious agents, and that illustrate a coordinating integration of body system dynamics. To arrive at this result, a molecular model is presented which shows, for the first time, that the hormone cortisol (CORT) and prostaglandin E2 (PGE2) have approximately equivalent chemical affinity, as indicated by the positioning of functional groups in hydrogen bonding and hydrophobicity, with the ligand binding domain of the glucocorticoid receptor (GR). A mathematical model is developed to predict that the signs and symptoms of illnesses are associated with the competitive inhibition at the GR of CORT and PGE2 within the hypothalamus that prevents normal gene expression during DNA transcription. To validate the pathways and model, a case study is presented to analyze the cause and presentation of fever and fatigue over multiple days due to the injection of a pneumococcal vaccine as influenced by physical activity. The research provides quantitative understanding of the root causes of signs and symptoms of infectious disease, which for example can offer a quantitative explanation of common symptomatic concerns of illness, such as fever, and can result in optimal drug treatment plans to minimize the effects of ailments.

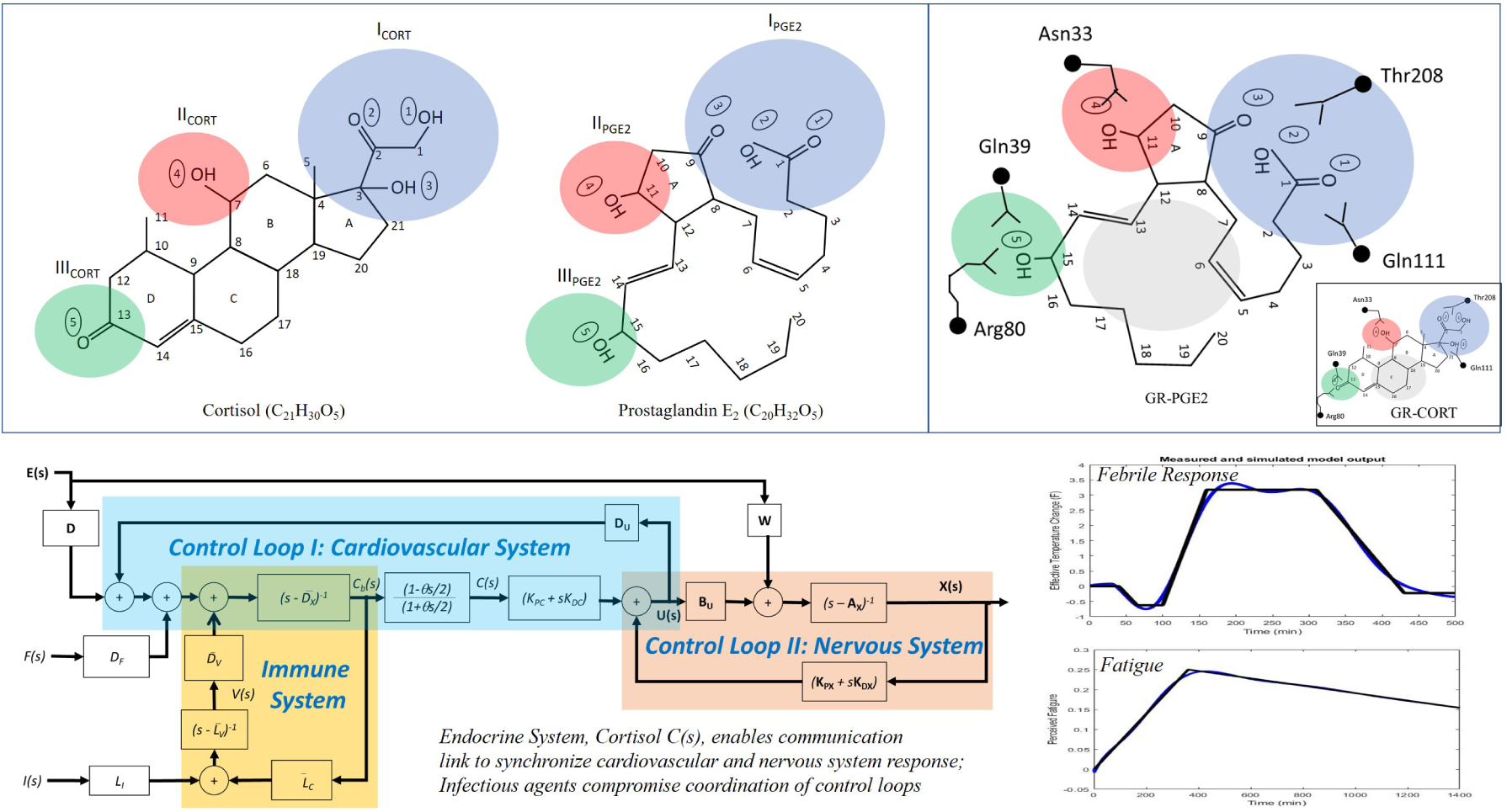
Graphical Abstract.

## 1 Introduction

While infectious disease, such as influenza and pneumonia, manifest in varied extent and duration, signs and symptoms of the associated autonomic dysfunction are actually quite similar across the human population, presenting as fever, fatigue, aches, congestion, brain fog, malaise, disruptive breathing and cardiac function. Although there are curative measures to reduce the impact of such illnesses, the fundamental origin for the decline in performance of the autonomic nervous system is unknown. Further, reasoning is lacking to explain the multitude of sequences and magnitudes observed of symptomatic responses, and even during nominal times without afflictions, the underlying basis of syncopation is unclear in terms of distributed autonomic control over the individual body systems, that is the mechanism of integration of the cardiovascular, nervous, endocrine and immune systems. This research describes a basic pathway coordinating the autonomic control system across body systems, and demonstrates how its dysfunction due to infectious agents results in observable signs and symptoms of illness.

The hypothalamus, located on the inferior portion of the diencephalon, contains a cluster of nuclei and tracts, and functions as the central controller influencing many body functions including those of the autonomic and endocrine systems [11]. Metabolism, heart rate, digestion, blood pressure, body temperature, limbic response, as well as stress-related factors, respiration [8], cerebral function, including memory [28], are all influenced by the hypothalamus. Sensory neurons terminate in the hypothalamus and in response efferent fibers extend into the brainstem and the spinal cord, where they synapse with neurons for example of the autonomic nervous system [25]. Notably, the hypothalamus connects to the endocrine system through for example the HPA axis (hypothalamic-pituitary-axis), which has received signifcant attention as characterizing the response of the body to stress and other factors [22, 9, 10, 4, 27, 14, 17]. The HPA axis involves the hypothalamic release of corticotropin-releasing factor (CRF) that binds to CRF receptors on the anterior pituitary gland, and in turn the release of adrenocorticotropic hormone (ACTH). ACTH binds to receptors on the adrenal cortex and stimulates adrenal release of cortisol. To complete the feedback loop, cortisol inhibits the hypothalamic release of CRF as well as the pituitary release of ACTH. Both cortisol (CORT) and its glucocorticoid receptor (GR) are widely distributed across cellular levels in which CORT, as the most abundant glucocorticoid, interacts in the cytoplasm with GR, and is then transcribed with DNA for protein expression [13]. The function of such corticosteroid action in the hypothalamus thus acts as a triggering mechanism [26] and synaptic potentiation have been noted [9]. In addition, CORT has a significant influence not only on the hypothalamus but on most bodily functions at the organ level, such as renal function [21].

To identify the pathway as it nominally functions, it is necessary to consider its performance when not working properly, for example due to the presence of pathogens capable of altered response. Perturbation to the normal homeostasis function enables an analysis, in this case molecular modeling and mathematical analysis, of the accuracy and predictive capability of the underlying theory. Of particular interest to this study is the correlation of CORT levels and symptoms similar to an infectious disease, even if the cause is not due to a pathogen. For example, dysfunction of too low cortisol levels due to misfunctioning adrenal gland can result in a crisis called adrenal insufficiency with symptoms that include fever, chills, fatigue, loss of appetite, stomach pain, body aches [18]. It is noted that these symptoms of adrenal insufficiency are similar to that of an infection, which is even at times misdiagnosed [5] as such, since it is rather due to cortisol levels - thus, it indicates a potential relation between cortisol levels and symptoms due to infectious disease. Further, there may even be a connection with Alzheimer disease, in which it has been noted that elevated levels of glucocorticoids and HPA axis dysfunction are consistently observed in patients [2]. Further correlations can be inferred from the undulation of fever characterizing disease such as brucella [6] and the rise and fall of CORT during the day and night [23] as well as respiratory systems during sleep [24, 1]. In addition, it is noted that injections of hydrocortisone can control a fever [3].

As motivation for this work, this research was initiated from an observed autonomic dysfunction of a symptomatic response involving fever, chills, and fatigue that was put in motion apparently by a vaccine injection of the pneumococcal agent followed several hours later by a regiment of physical activity. Current theories of thermoregulation would indicate that the reason for the fever was an adjustment due to the presence of the bacterial agent that was made to the temperature “setpoint” located in the hypothalamus that the body functions subsequently responded automatically to accommodate the new higher setpoint resulting in the observable temperature trajectory of fever. However, that justification seemed to this investigation to be inadequate, for example it would suggest that it was mere coincidence that the altered temperature response began approximately fifteen minutes after the physical activity was completed, which motivated the presently described research. To analyze the observed responses, the initial direction of research involved the development of a mathematical model to describe the temperature dynamics in terms not only of the nervous system but also the cardiovascular system, which was needed in order to integrate the influence of physical activity on the resultant response of chills and fever [20]. While the general structure of the proposed mathematical equations of the underlying theory describing the temperature response were adequate to explain the observed trajectory, the molecule that linked the cardiovascular and nervous system was not identified, but was referred to as a mediator and reporter involved in an association with the infectious agent. Here, this linking molecule is identified that modulates performance of the autonomic nervous system, and whose dysfunction results in observed autonomic-related signs and symptoms of infectious disease.

The results comprise a mechanism involving a molecular model that *for the first time* shows the ligand binding domain of the glucocorticoid (GR) receptor has approximately equivalent affinity for the hormone cortisol (CORT) as it does for prostaglandin E2 (PGE2), which is present with certain infections and associated with the inflammatory response. A mathematical model is developed that predicts symptomatic responses, such as fever and fatigue, as a reduction in performance due to the competitive inhibitory interaction of GR with PGE2 instead of CORT which thereby prevents nominal gene expression by transcription of DNA. The mathematical model is then analyzed in its capability to predict the symptomatic responses of body temperature and fatigue due to the vaccine and subsequent physical disturbance. The application of this research is then described for the diagnosis of ailments and to minimize symptoms of infectious disease with optimal behavioral and drug treatments. It is shown that dysfunctional alterations to just this single hormone cortisol can explain the multitude of signs and symptoms of illnesses, whose sequence and extent is a function of the external load, the trajectory of the infectious agent concentration, and the sensitivity function of the internal autonomic control system.

## 2 Results

In this section, results are presented to indicate pathways of autonomic control function and dysfunction, of which the molecular model that shows for the first time the equivalency of CORT and PGE2 with GR, thus creating competitive inhibition of CORT within the hypothalamus in the presence of infectious disease and inflammatory conditions. To further address this issue, a mathematical model is developed that links the cardiovascular, nervous, immune and endocrine systems to evaluate the effect of such competitive inhibition of GR created through interaction of CORT and PGE2. The predictive capability of the model is evaluated in a case study on fever and fatigue due to an adverse response to pneumococcal vaccination.

### 2.1 Molecular affinity of CORT and PGE2 with the LBD of GR

In addition to functionality during normal conditions, of particular interest is the pathway and mechanism by which cortisol is impacted by infectious agents. Here, PGE2 is assessed as it is commonly associated with illness in response to bacteria which contain lipopolysaccharide (LPS). PGE2 is also generated a through a reactive sequence results involving arachidonic acid (AA) and cyclooxygenase-2 (COX2), among other mechanisms including interleukin-6 (IL-6) [7].

In Figure 1, the chemical structure of CORT and PGE2 is presented in the standard configuration, in which a striking similarity is noted. CORT and PGE2 are of similar molecular weight, 362.46 g/mol versus 352.46 g/mol respectively, and have nearly the same number of chemical elements 21 versus 20 carbon; 32 versus 30 hydrogen; and importantly the same number of oxygen elements, including carboxylic, hydroxyl, and carbonyl groups. CORT has a five membered ring as does PGE2 which can be formed notably from interaction of AA and COX2.

**Figure 1:**
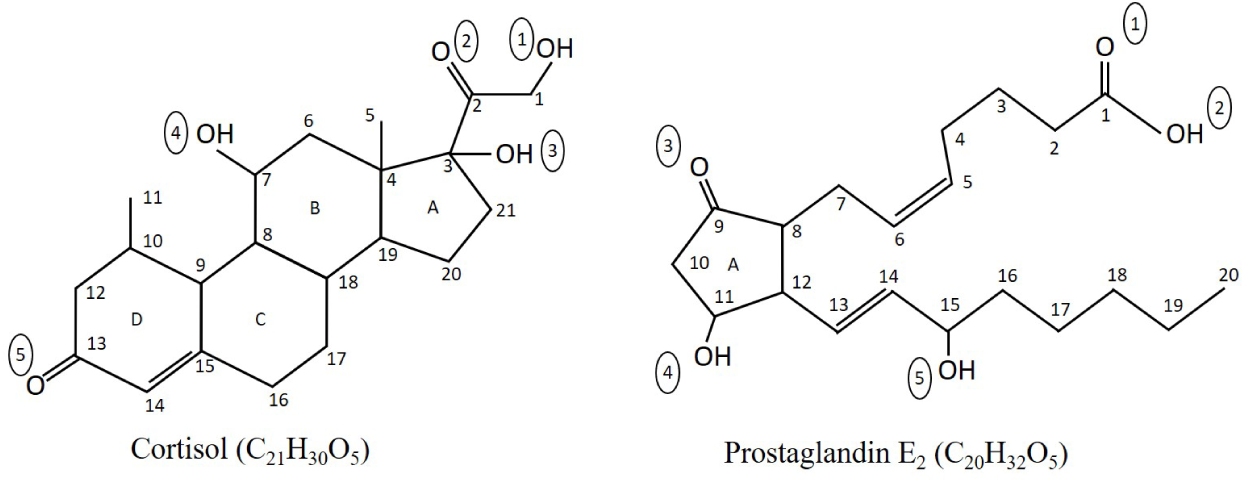
Expressed in standard format, a comparison of cortisol and prostaglandin E_2_ molecules to denote the similarity in the number of chemical elements of carbon, hydrogen, and oxygen, hence molecular weight. Moreover, the number of oxygen related functional groups is five, and is equivalent between the two molecules with 3 hydroxyl groups including one at the end, and 2 carbonyl groups.

When expressed in standard format the structure of PGE2 is dissimilar to the ring structure of CORT; however through a rotation at C4 and C13 of PGE2, a structure resembling a set of rings is formed as indicated in Figure 2. This expression maintains the same double bond arrangement of *cis* configuration on C5-C6 and *trans* configuration on C13-C14. It is also noted that the arrangement critically positions three oxygen groups together such that it is comparable in position to that of CORT.

**Figure 2:**
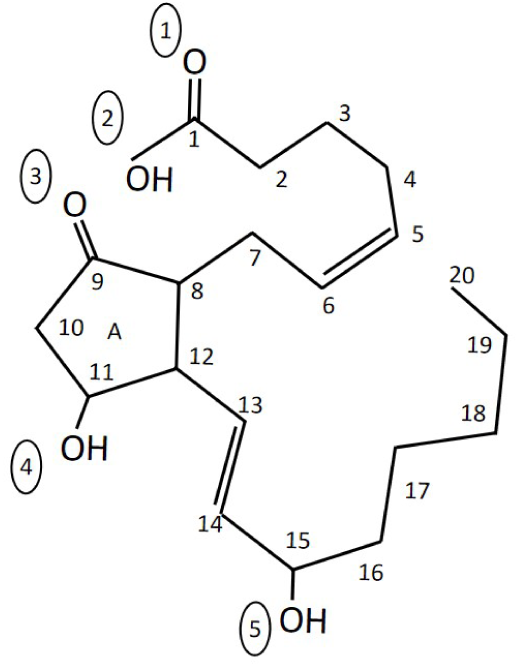
To establish the positioning of PGE2 to better match the activity of cortisol, the structure is proposed where there is a repositioning of PGE2 at C4 and C13. The upper branch, relative to the five membered carbon ring, is positioned such that three oxygen groups are in close proximity and the carbon groups form relative ring structures. The bottom branch is rotated such that a ring formation is roughly formed and the two double bond groups are placed in a linear path, while moving the hydroxyl group along the outer boundary of the structure. Note that the C5-C6 cis and the C13-C14 trans double bond structures are maintained.

In Figure 3, the functional groups between CORT and PGE2, with rotation, are compared to demonstrate approximate chemical equivalency. In the group of I_CORT_ and I_PGE2_, three oxygen groups are present, and with II_CORT_ and III_CORT_, II_PGE2_ and III_PGE2_, one oxygen group each. At the interior of the arrangement, a hydrophobic center exists, with the two double bonds well positioned in PGE2 to duplicate the ring structure of CORT. The greater motion of the carboxylic group of I_PGE2_ is available to offset the one carbon offset of the equivalent structure of I_CORT_. The similarity is also present in the relative positioning of the oxygen-related functional groups of the two molecules.

**Figure 3:**
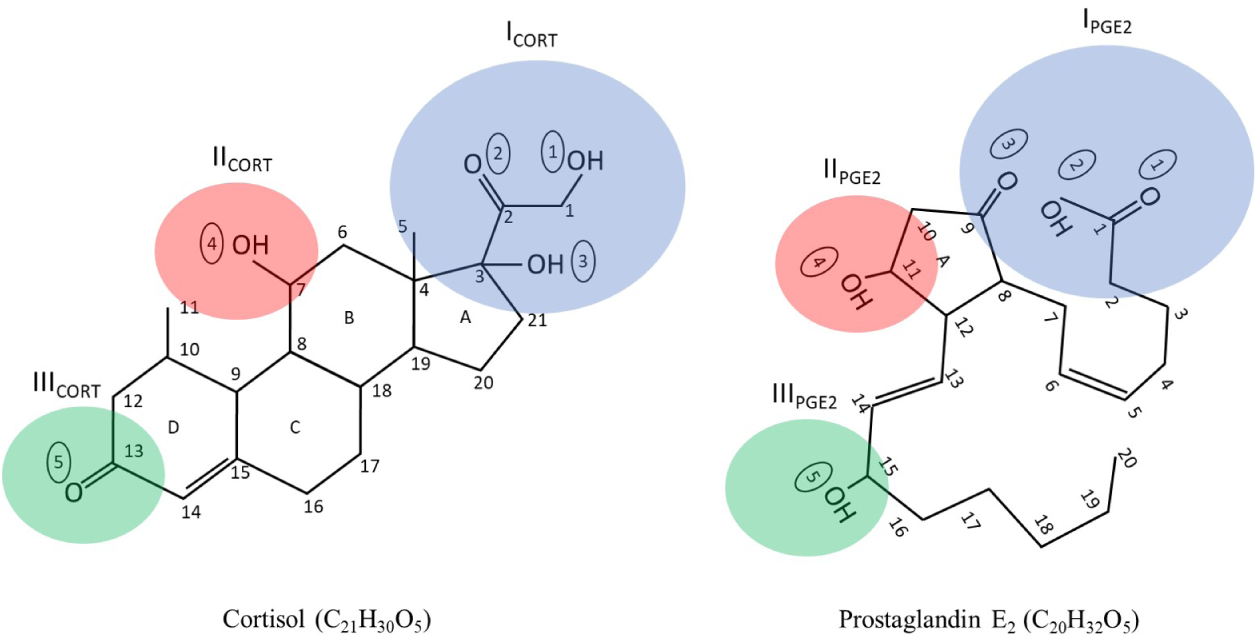
With the reconfiguration of PGE2 and with a rotation of approximately 45 degrees, the cortisol and PGE2 molecules are compared for the functional groups and relative positioning. It is noted that the three sites are correlated with I_*cort*_ and I_*cort*_ having three oxygen groups, with one each on the other two groups. The positioning of the functional groups is on the exterior of the PGE2 molecule, equivalent to cortisol, and the interior is composed solely of carbon and hydrogen groups.

Based on recent high resolution crystal structures of the interaction of CORT with the ligand binding domain (LBD) of GR [15], the approximate positional relationship of PGE2 with GR, instead of CORT, is presented in Figure 4. It appears to be a nearly identical fit between GR-PGE2 and GR-CORT, although apparently it will function dissimilar and not result in DNA expression in the same manner of CORT. It is noted that there is significant hydrogen bonding at the three functional sites, and may even be preferred in comparison to CORT as III_CORT_ requires the presence of water, whereas with III_PGE2_ the hydroxyl group is available to interact directly with Gln39 and Arg80. In I_PGE2_ the single hydroxyl group is available for interaction with Gln111 and Thr208.

**Figure 4:**
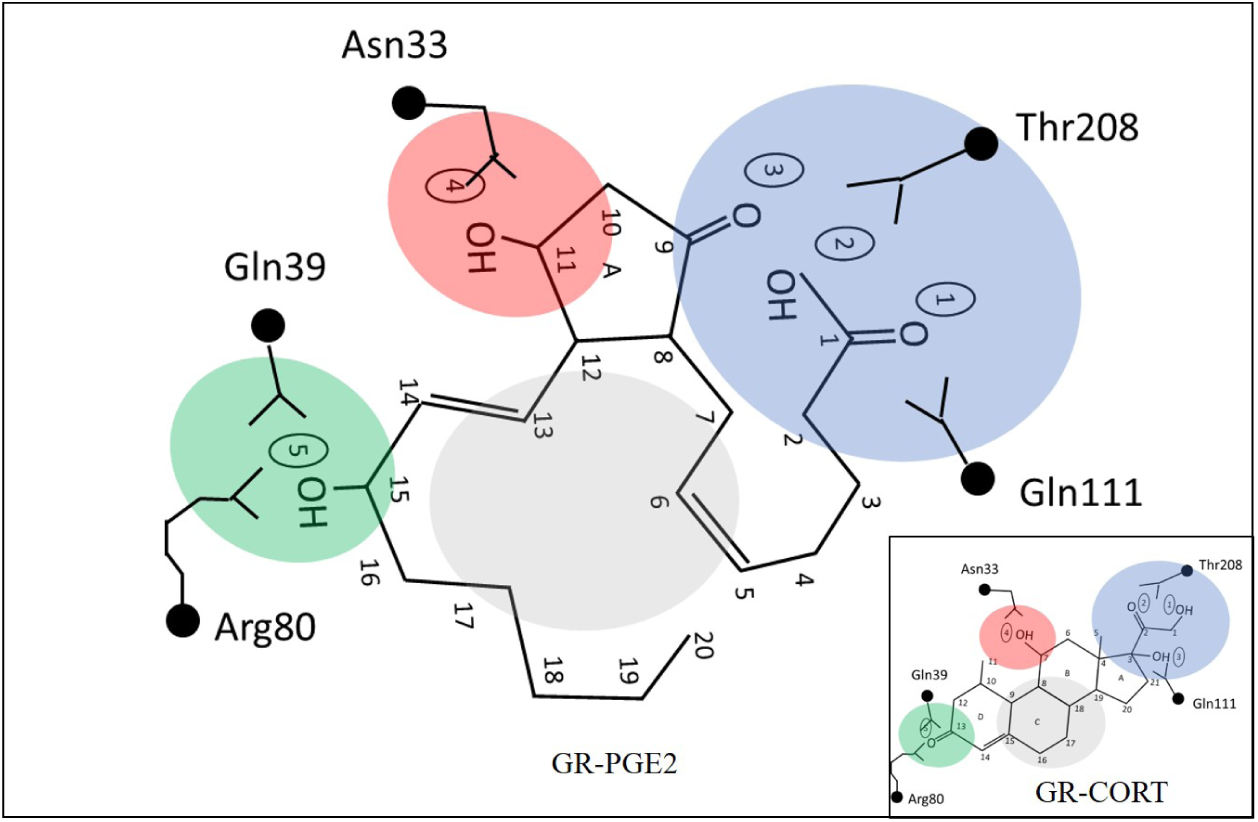
For the ancestral glucocorticoid protein receptor (AncGR2), which is a stable GR useful for crystal studies, at the ligand binding domain (LBD), a fit of PGE2 very similar is noted to that which is achieved with CORT as indicated in the inset. There is substantial equivalent hydrogen bonding at the three functional groups, and at the interior there is substantial hydrophobic interaction and similarity. There may be an improved interaction with PGE2 relative to cortisol with Gln39 and Arg80 (green shaded region) since CORT requires an external water molecule since only a carbonyl group is available, whereas PGE2 already has a hydroxyl group. The three membered oxygen group of PGE2 contains enough free motion as it is unconstrained by rings to achieve optimal alignment (blue shaded region). The double bond alignment of PGE2 at the hydrophobic core is noted. It is also noted that the ring structure of PGE2 formed via the C8-C12 bond is critical to achieve the proper configuration of the two branches and subsequent fit within the hydrophobic core (gray shaded region), which incorporates two carbon-carbon double bonds. Thus the effectiveness of NSAIDs, which disrupts the formation of the C8-C12 bond, is apparent.

Further examination of Figure 4 also indicates the importance of the five-membered carbon ring formed by COX2, as it positions the two branches of PGE2 to associate with the hydrogen bonding centers of LBD of GR. It is noted that the use of aspirin and NSAIDs to block COX2 and thus the suppression of PGE2 would result in improvements of symptoms such as fever.

Moreover, the ring structure of PGE2 sets-up the functional groups on the exterior of the molecular and an internal hydrophobic core, which includes a pair of carbon-carbon double bonds at C5-C6 and C13-C14 interacts with groups of GR including Leu77, Met73, Trp69, Met70, Leu32, Phe92, Cys205, Phe204 (not shown in figure but positioned below the internal core of PGE2). The internal hydrophobic core for each of CORT and PGE2 includes six carbons.

We believe that these modeling results are the first indicating the interaction of PGE2 with GR, displacing CORT. Importantly, it indicates competitive inhibition at the ligand binding allosteric site of GR. It can be inferred that other structures where CORT interacts, such as membrane bound receptors that produce non-genomic responses, that there would be similar competitive inhibitory influence of PGE2 on CORT.

### 2.2 System Modeling of Cortisol-Hypothalamic Pathways including Dysfunction

To develop a model of the influence of CORT, and the competitive inhibition of PGE2 at GR, a mathematical model of body systems integration is presented in which signs and symptoms of illness can be assessed. While such signs and symptoms pertaining to the hypothalamic response associated with infectious disease can be listed, the severity and sequence of each is variant. To better quantify this effect, and to validate the hyptothesis put forth of CORT dysfunction as the source of abnormal body system responses, the model is used to assess why some symptoms are moderate and some severe; Why sometimes there is fatigue with fever and sometimes only fatigue without fever. Although it would seem to be of a multivariable nature, in this section, we show how the variation of just a single hormone, CORT, is explanatory. The resultant model is a lumped parameter structure that can be identified from input-output data and validated by its analysis and predictive capability, as well as its internal structure An important aspect of this analysis is the feedback control law postulated for the hypothalamus, as it will determine its overall integration and coordination within the network of systems, and its interaction with causal factors, including infectious agents and internal stimulation. From a high level, the model is a neural and neuroendocrine mapping of the inputs and outputs, as indicated in the schematic of Figure 5.

**Figure 5:**
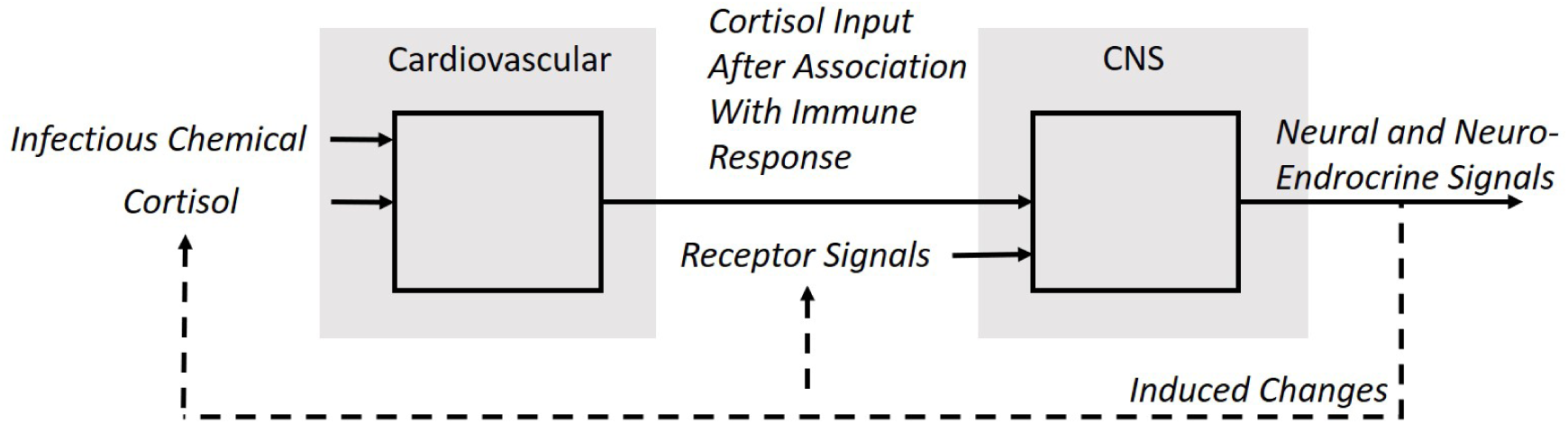
A simplified schematic of the control system is depicted as follows starting from the left: cortisol and the infectious agent interact in the sense that the infectious agent associates with the receptors that nominally CORT would utilize prior to a genomic response, which is considered as within the cardiovascular system; thereby alters the concentration of the CORT from its nominal expressive value during transcription. The CORT level that does makes its way through the plasma membrane generates neural and neuroendocrine signals associated with the receptors to generate signals associated with system effectors, such as body temperature regulators, breathing rate, heart rate, and other basic autonomic functions. These autonomic signals then result in changes to the body conditions, which manifest in changes to both the receptors and to cortisol concentration by the manipulations that caused the adjustments in to be made.

#### 2.2.1 Feedback Control

To quantify the dynamics and control as dictated by a newly proposed cortisol-hypothalmic mechanism to explain the symptoms of illnesses associated with the autonomic nervous system, a mechanism is developed by which the internal commands are generated from neurological receptor signals in combination with CORT, which establishes physical status of the cardiovascular system. The control law is postulated as s a function of the levels and its rate of change associated with the incoming receptor signals and the concentration of the cortisol along with its rate of change, as follows:

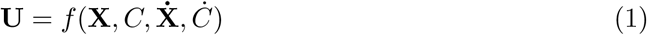

where **U** indicates the neural and neuroendocrine signals from the hypothalamus to control around nominal levels, **X** denotes the incoming signals associated with the body status such as temperature, blood pressure, *C* is the chemical signal indicates cortisol concentration, 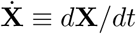 and *Ċ* ≡ *dC/dt*, which are the rates of change associated with the body status signals and cortisol respectively.

The rationale for incorporating the rate of change as a signal of the internal controller, which is first described in earlier work of this lab [19], is that the concentration gradient at the synaptic cleft or within the cytoplasm will define the speed and amplitude of the internal responses, such as temperature and fatigue. The concentration gradient is associated with an open system and subject to second order partial differential equations of a flow fields. The boundary conditions associated with such models take into account the spatial gradient, and the time derivative associated with the input of *C* and signals **X** will establish the spatial concentration gradient. In [20], further discussion is given for the use of the rate of change in developing a control law.

A schematic of the control law, which is determined associated with the central nervous system, is depicted in Figure 6 in which the linkage with the cardiovascular system is also shown for clarity, including the infectious agent *V* and CORT *C*_*b*_ prior to its interaction with GR.

**Figure 6:**
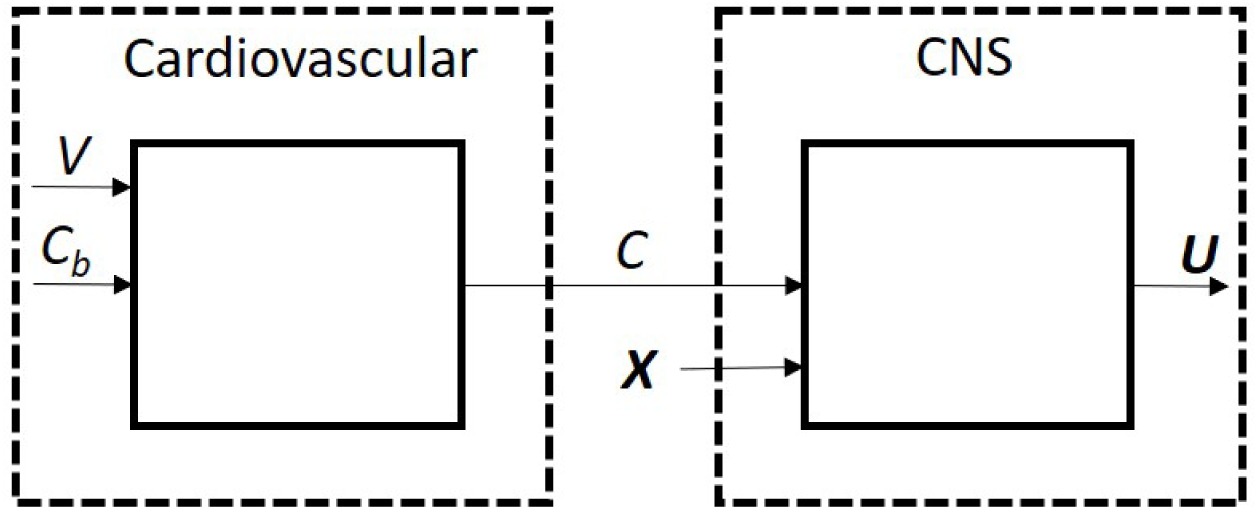
A schematic of the control law focuses on the input-output relation of the central nervous system (CNS), but it is noted for completeness that the signal entering the CNS, C, represents the concentration of cortisol. The infectious agent V is correlated to the concentration of prostaglandin E2, which interferes with the transmission and action of cortisol through its association with the glucocorticoid receptor. The autonomic responses are represented as a neural signal **X** and the output from the CNS is U which are neural and neuroendocrine signals. Cortisol C can cross between the blood-brain barrier in order to produce such changes in conjunction with the receptor signals, thereby linking the cardiovascular and nervous system in terms of the internal controls associated with the hypothalamus. The output signal undergoes further processing to produce effector signals at the target sites.

The perceived or effective input signal **X** is one that need not be explicitly measured, but rather a signal that is representative of the symptom or sign, such as fatigue or body temperature. The output of the manipulated neural signal, **U**, may signify a combination of responses to the system, such as increase or decrease in vasodilation, sweating, shivering, respiration rate, and other such mechanisms employed for automatic control as programmed through the hypothalmus. For the purposes of this study, the distribution of the output signal to the effector mechanisms is secondary to the analysis and can be taken-up for study as it will not impact the present analysis. Thus, to obtain a mathematical representation of the control law, a Taylor Series expansion of Equation (1) is taken about a nominal operating point denoted as **U**_0_, **X**_0_, *C*_0_, which would be associated with a standard body conditions, and truncated after the first term results in a linear expression. Deviation variables are defined as Δ**U** = **U** − **U**_0_, Δ**X** = **X** − **X**_0_, Δ*C* = *C* − *C*_0_, and thus we have the control law expressed as:

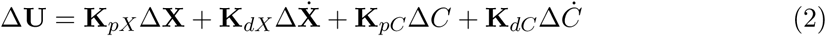

where the gain functions denote the sensitivity about the operating point as discussed in the supplementary material of [20]. It is noted that the control law, Equation (2), is considered as having a PD (proportional-derivative) form, which is deployed in industrial process control with success. The proportional terms provide a means of using a feedback control mechanism to manipulate the input variable so as to steer the controlled signals about a nominal value. The derivative terms is primarily to shape the response to desirable trajectories, for example speeding-up the control action, but does not in general influence the stability of the system. Industrial controllers have benefitted from an integrating action, in which the error in the control signal is driven ultimately to zero; however it does introduce instability and there is no evidence that an integrating action is incorporated. It is noted that this analysis incorporates a feedback mechanism that is based upon a chemical signal that indicates the status of the input in response to the output signal. Moreover, based on observable responses to infectious agents, it appears mandatory to incorporate such a coupling mechanism, since without it, and a feedback controller that just fed back on the status of the output signal **X**, and not the input signal, would not be able to capture the impact of infectious agents.

#### 2.2.2 Body System Dynamics

The body system signals, which are a combination of neural and neuroendocrine inputs, by which control signals are generated according to the control law of Equation (2) corresponds to an effective signal that is expressed as a function comprised of not only body system variables **X** and *U*, but also exogeneous inputs, [*E*_1_, *E*_2_, …, *E*_*n*_] that can have effect on body systems, such as exercise, environmental factors like external temperature, and other inputs including stressors such as those associated with deviations of the sympathetic and parasympathetic nervous system. Thus *E*_*j*_(*s*) can incorporate inputs due to emotional stress, cerebral thoughts that may induce sympathetic activity, and other activities associated with parasympathetic activity. In the approach developed in this paper, the representation for body system dynamics is developed in closed-loop format for general sets of inputs. As the body system responses are in general non-specific, but associated with perception, the signal of concern here is considered as “effective” or “perceived”, that is one in which the control laws ultimately respond after integration of the neurological receptors and signals. The effective system dynamics are expressed as a linear dynamic model obtained from a Taylor Series expansion and then neglecting all terms of order two and higher, and in utilizing deviation variable format with Δ*E*_*j*_ = *E*_*j*_ − *E*_0,*j*_ about a nominal level, which could be 0, an expression for the body system dynamics, such as for temperature, fatigue, is achieved:

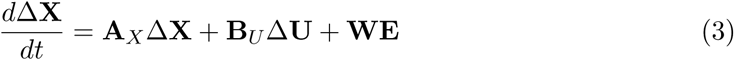

Equation (3) is a linearized model with system matrices expressed as sensitivities, see the supplementary material [20], which maps the body state trajectories, inputs from the control law, and exogeneous inputs to changes in the body system outputs, such as temperature and fatigue, and is typical of the dynamic response of physical systems when used for control system analysis about nominal operating windows.

#### 2.2.3 Cortisol Dynamics

The development of the feedback control law for the hypothalamus, it is proposed that cortisol is the critical co-controlling factor, as it is indicative of the status of the effects of the controller output signal in its response to body system variations. It is further developed that cortisol is influenced through infectious agents through competitive inhibition with GR, and thus a rate of association is defined as *R*, which is the extent by which the infectious agent interferes with the control signal through coupling with the glucocorticoid receptor.

Further the external inputs to the system are considered to be potentially influential on CORT dynamics, such as the effects of exercise. Also incorporated is the potential for an external feed stream *F* that can provide an external source of CORT, such as hydrocortisone injection. The resulting function is expressed therefore as follows:

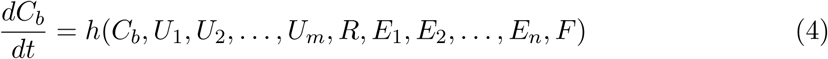

and in producing a linearized representation of Equation (4), a Taylor Series expansion is truncated after the first term. For the reaction rate *R*, we consider a reaction law which is dependent upon the product of the concentration of the external reactive component *V*, such as a bacterial or viral infective species, which would influence the concentration of PGE2. The reaction rate would also be a function of CORT. For competitive inhibition, a nonlinear relation following for example Michaelis-Menten kinetics would seem appropriate, and here the reaction rate is selected in genaral terms as:

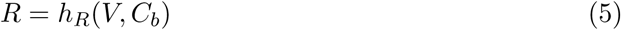

While the rate coefficients may be a function of temperature or other body system states, it is considered that such effects will be secondary and thus can be neglected without loss of generality. Linearizing the reaction rate representation of Equation (5)yields an overall relation for the concentration of CORT:

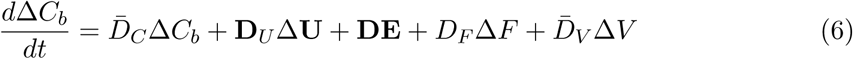

where

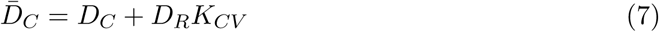

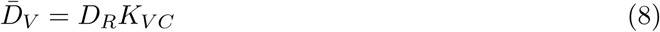

Note that the dynamics of CORT, 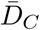 are influenced by the effective overall reaction kinetics *K*_*CV*_ of the external agent, *V* and thus PGE2 interacting with GR and hence impacting the effectiveness of the autonomic control system as driven by CORT.

#### 2.2.4 Infectious agent dynamic model

For the infectious agent *V*, its dynamic model is represented from a species balance that includes *I* as a mechanism by which the invasive agent enters the cardiovascular system, such as with an injection which can include an internal developmental period subsequent to release into the system. A Taylor Series expansion truncated after the first term yields after substituting for Equation (5) *R* and taking deviation variables, Δ*I* = *I* − *I*_0_, yields

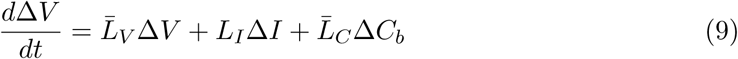

where

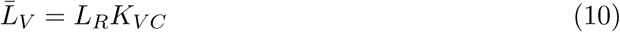

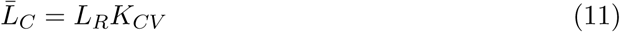

#### 2.2.5 State-space model

To form the system response of Equations (3), (6),(9), the state variables are defined as **ΔY** = [Δ**X**^*T*^Δ*C*_*b*_Δ*V*]^*T*^ and the disturbance variables as Δ**Z** = [Δ**E**Δ*F*Δ*I*]^*T*^ which can be related in state-space representation

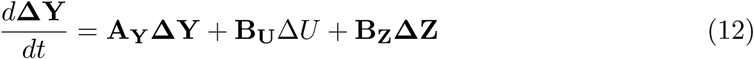

with the system matrices,

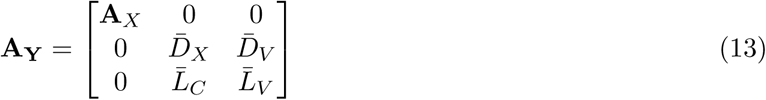

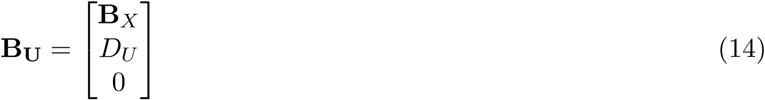

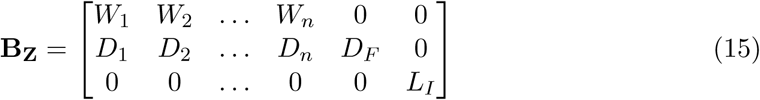

This state-space form has the general solution [12],

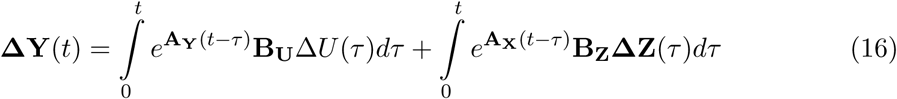

The response, depending upon the nature of the system matrices, is in general comprised of a series of attenuated trigonometric oscillating functions, which in general does exhibit consistency with observed responses, for example a oscillating temperature response expressed as chills and fever, hypothermia and hyperthermia.

#### 2.2.6 Time Delay

The location of the control response to the concentration of CORT may be distinct relative to that of the local anatomical sectors responsible for its generation, or for the expression of the response from CORT such as its interaction via GR with DNA, and thus the transportation delay is modeled by a time delay function:

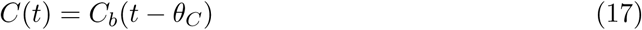

where *θ*_*C*_ denotes the time delay to gene expression.

#### 2.2.7 Dynamic System Response

In forming a set of linear, time-invariant models to describe the closed-loop body system state dynamics, the LaPlace transform, denoted as *s*, of Equations (2), (3), (6),(9) is taken. The closed-loop schematic representation is presented in the block diagram of Figure 7, which maps the inputs to the system *E*(*s*), *I*(*s*), *F* (*s*) to the output body system signals *X*(*s*). The internal control loop is highlighted indicating the intersection of two coordinated control loops. To arrive at the final set of equations to describe the dynamic response and internal control operations of body systems, the model relations are linear approximations of the functionals describing system performance and that the nonlinear behavior of the system is characterized by a sequence of linear, time-invariant models taken around nominal operating points.

**Figure 7:**
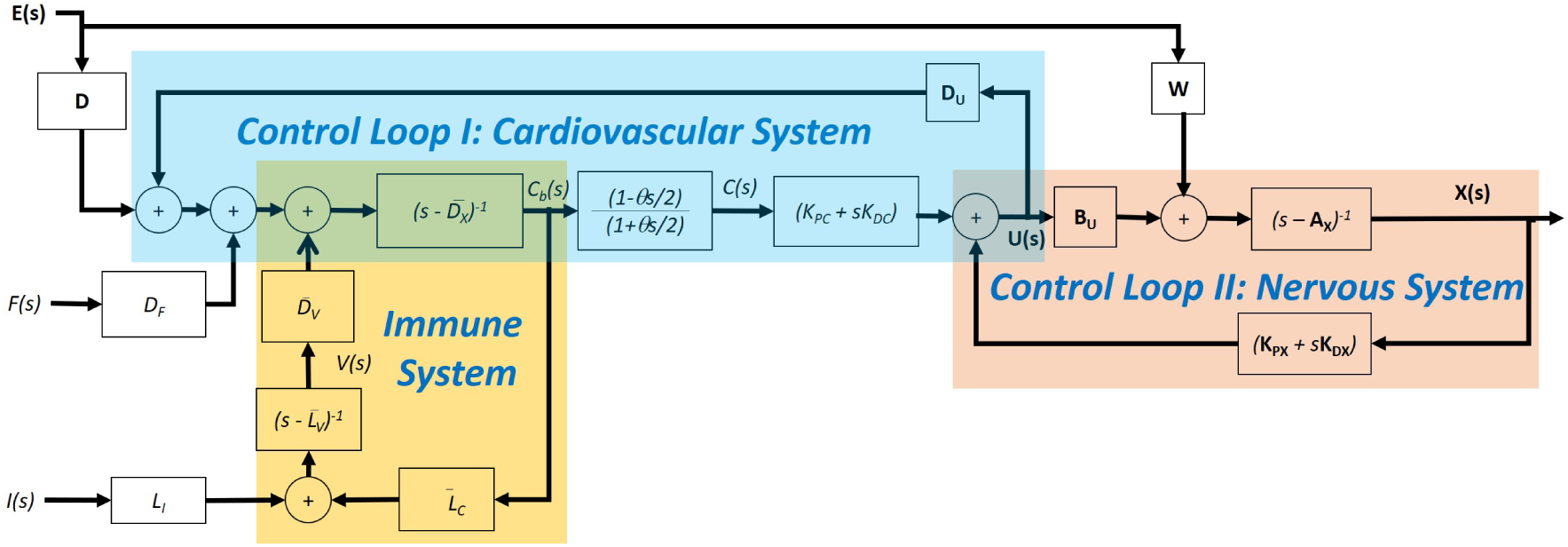
The block diagram of the closed-loop response of autonomic system functions in response to inputs from infectious agents V (s), from external agents F (s), and from a vector of external sources, **E**(s), which comprise external stressors. The blocks comprise the transfer functions of the model, and include the control law function for **U**(s) which comprises feedback from body state dynamics **X**(s) and cortisol C(s). The schematic of the feedback control law and model is partitioned to indicate interactive and cooperative control loops of the cardiovascular system and the nervous system, as well as the inputs via the immune system into the cardiovascular system. Note the cooperation of the cardiovascular and nervous system, as for example, the cardiovascular system responds to commands, for example from the need for increased cellular oxygen due to sympathetic stimulation, the nervous system will respond automatically by feedback control to regulate autonomic system functions, thereby maintaining a homeostatic internal environment. The converse in cooperative feedback loops holds as well, for example with external disturbances, due for example to an increase in external environmental temperature, are automatically compensated by the feedback controller of the cardiovascular system through the control law of Equation 2.

A interesting consequence of the proposed control law mechanism is an intersection of signals associated with the cardiovascular system and with the nervous system linked through the hormone CORT, and includes the immune system response which is in competitive inhibition with CORT at the GR within both the hypothalamus and target organs. The intersection is indicated in Figure 7 with the disturbances induced from external sources, **E**(*s*). The inputs to the control law, whose location is the hypothalamus region, are chemical and electrical in nature, and the output of the control law is neurological and neuroendocrine. Note that the block diagram, which compartmentalizes the immune response with the cardiovascular and nervous system, is formulated as a graphical representation of the mathematical equations describing the process. The control loops are cooperative in the sense that stressors placed on the cardiovascular system, such as increased oxygen demand, can be balanced by the requisite body system response, such as adjustments in body temperature. And conversely, stressors placed on the nervous system, such as the need to dissipate heat, can be balanced by the circulatory response, such as with vasomotor activity. Conversely, the control loops are not run independently, in that changes made to the cardiovascular system are conducted in conjunction with regulation of the nervous system and thus other body function systems.

### 2.3 Case Study

To test the validity of CORT and PGE2 exhibiting competitive inhibition with GR, and to test the validity of CORT as the hormone in control of the symptomatic responses of illness, and to validate the mathematical model, the following case is analyzed: An administration of a subject by injection of a vaccine associated with pneumococcal species was given at approximately noon. The temperature was constant prior and immediately after the vaccination, and throughout a two-hour period of exercise that took place six hours after the injections. Approximately ten minutes after completing the exercise period, which consisted of moderate weight training and a seven-mile jog on a treadmill, the sensation of severe chills was felt that lasted approximately forty minutes followed by a rapid rise in temperature to a nominal fever, which lasted two hours, followed by sweats in a return to a baseline temperature, thereby concluding the episode seven hours after the completion of the exercise period. No reports of distress were reported after the episode of temperature fluctuation. On Day 2 and on Day 3, following the same exercise protocol at the same time of day, a similar response was reported in temperature response, through much less severe, of which the intensity of the response was perceived as level 8, 3, and 1 for each of the three days, and the period of response was longer during the chill period but shorter in time at the higher temperatures. The first day required bed rest during the seven hours of high temperature excursion to deal with the chills and fever, while the second and third day did not require bed rest. By the fourth day, no adverse temperature effects were reported following the exercise protocol of a seven mile jog. It was noted that for the pneumococcal vaccine, similar experiences of chills followed by fever were reported from on-line patient feedback, hence the display of temperature effects following vaccination was deemed to be fairly typical, although this case was particularly well characterized as the exercise regimen was a precursor to a significant dynamic temperature response The reports were that the exercise protocol was normally 9 miles but could only sustain 7 miles before the onset of fatigue. For the second day of exercise, fatigue was reported as half as significant as the first day.

To evaluate the ability of the model to produce the characteristic temperature response, a model for each of the three days is determined using system identification techniques [16] available in the Matlab programming language with the forcing function of a step-down in *E*_1_(*s*) considered as the exogenous input of exercise activity. As consistent with the model derivation, the model order was selected as four poles and three zeros, and the mean square error was minimized in determining the model parameters. Using the multiple, linear time-invariant model representation about each new operating point, a model was produced for each of the three days. The temperature response is compared with the data in Figure 8, which show agreement in mapping the inverse response corresponding to the hypothermal condition of chills, followed by the temperature rise, either plateauing or momentarily hitting a peak, and then terminating by reduction to a baseline temperature.

**Figure 8:**
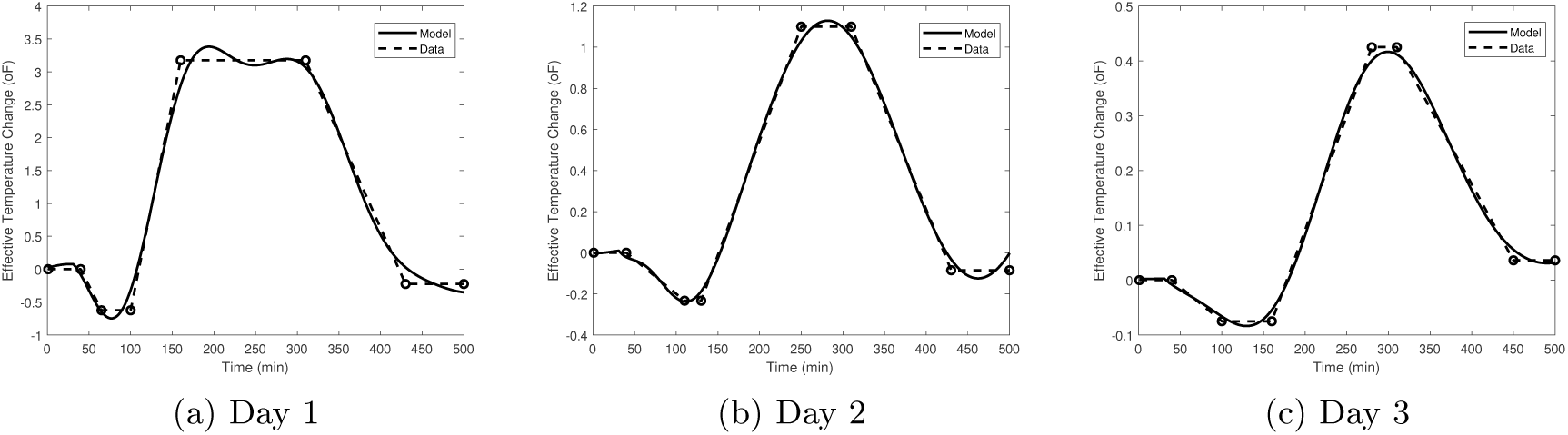
As an initial indication of model validation, a comparison of the measured and modeled trajectory of perceived temperature is compared for each of the three days. The inverse dynamics of the model are clearly captured in showing a dip down in perceived temperature followed by a rise and then a cool down back to base. For Day 1, some oscillation at the level off period in temperature is noted, while the static gains in settling the response for each of the days is indicated. Thus, the relatively close representation of the model output to that observed would indicate that the modeling approach is appropriate and that it can be examined for predictive and analysis capability.

A model for the fatigue response is also generated by considering that after the injection, the fatigue factor was perceived and was set at 25% reduction from nominal for the first day and 15% fatigue factor for the second day, and no fatigue reported after four days. A model was established as four poles and three zeros representation and computed from the input and response data. A comparison of the results for fatigue is presented in Figure 10 to show model prediction.

**Figure 9:**
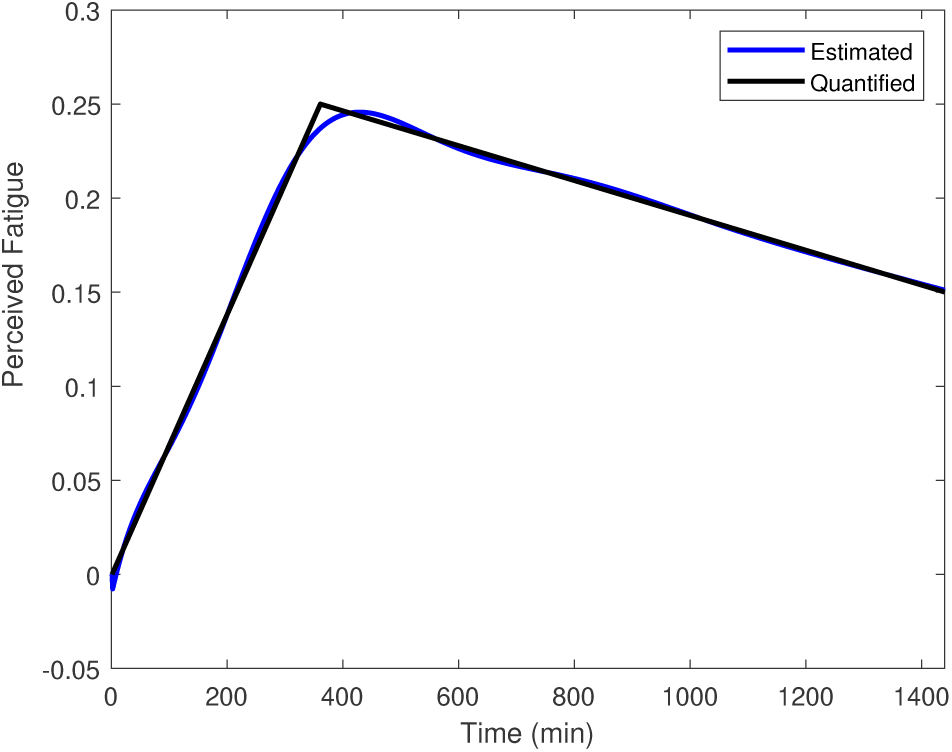
A comparison of the perceived fatigue factor over a two day period is compared with the model trajectory, which is seen to capture the slow transient response peaking after the first day and then subsiding. The model is of the order of four poles and three zeros with input excitation of the injected vaccine.

**Figure 10:**
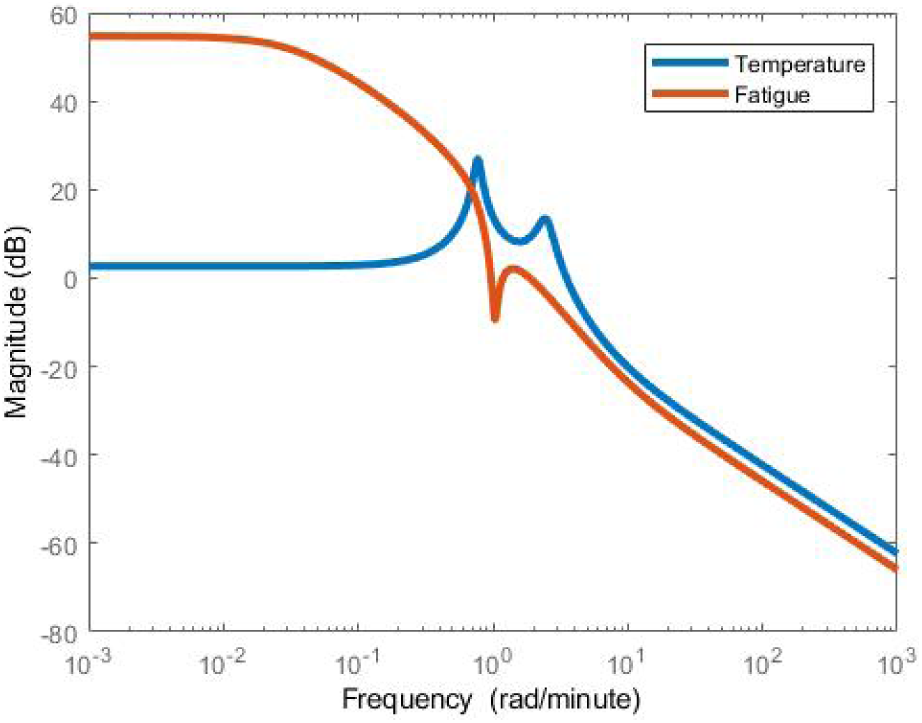
The magnitude of the frequency response of the models characterizing perceived body temperature and fatigue. The responses indicate that fatigue will be a slow effect and fast variations to the body will go largely attenuated, whereas body temperature adjusts well to slow variations, but is sensitive at certain higher frequencies loads, as indicated by the resonant peak. This is indicative that the same causative factor, that is the interference of nominal cortisol response due to the presence of the infectious agent uptake with the GR, can produce greatly different responses in this multi-output, single-input system.

The frequency response comparison of temperature and fatigue responses is presented in Figure 10 to denote the spectral separation in the responses, as well as the resonant frequency for temperature. This indicates the method in which responses are felt by the body systems: For example, temperature has tight control and will remove slowly varying loads as represented as low frequency loads, whereas fatigue will be felt by slow variations in load disturbances. However, fast distribuances will largely be attenuated for fatigue, such as rapid blood pressure and respiratory rate changes, although long term, slowly varying changes will influence fatigue.

The Nyquist plot of the frequency response is presented in Figure 11 in comparing the dynamics of Day 1, Day 2, and Day 3, as the body transitions higher concentrations of the vaccine to lower concentrations as it is depleted. The vaccine is seen to have a significant effect on the dynamic response, and based on the Nyquist stability criteria involving the encirclement of the point (−1, 0), would result in an unstable system if utilized with an additional layer of feedback control. Clearly, while infected, the temperature dynamics are significantly compromised.

**Figure 11:**
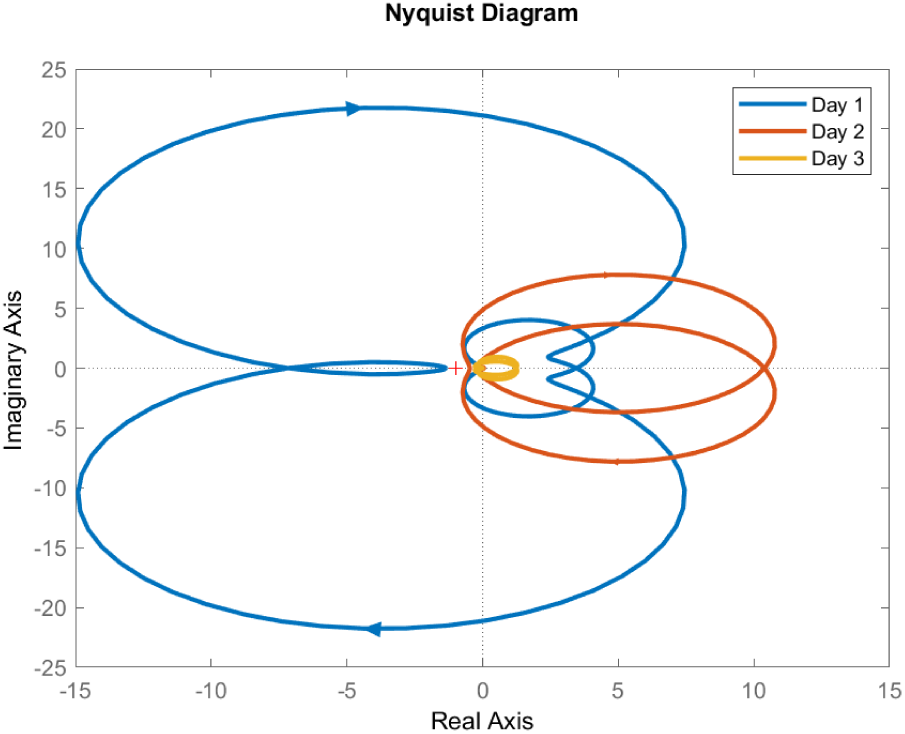
In evaluating the impact of the vaccine on body temperature control, the Nyquist representation of the models for Day 1, 2, and 3 is compared to show dramatically the severe degradation of temperature control due to the influence of the polysaccharide on the internal dynamics. The Nyquist plot shown an encirclement of the (−1, 0), which indicates instability if a secondary feedback controller were to be wrapped around the intrinsic closed-loop temperature control system. There is a significant reduction in the representation in Day 2 and in returning close to nominal behavior of Day 3. The result shows that infectious agents strongly affect temperature control system in putting the body at risk for instability if not handled appropriately.

To apply the model, the amount of CORT that could be taken externally, that is through the external feed stream *F* (*s*), to counteract the feverish response can be determined hypothetically through an estimate of CORT concentration change during fever according to the model response. It is necessary to estimate the parameters of the control law of Equation (2), which can be expressed equivalentally as Equation (18),

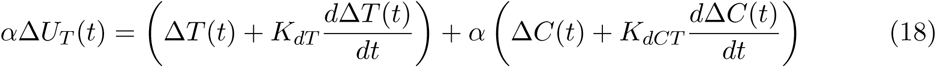

where *α* = 1*/K*_*pT*_, *K*_*dT*_ = 100, *K*_*pCT*_ = 1 and the derivative term of the concentration, *K*_*dCT*_ was compared for CORT levels using the values of 100 and 10, with the scaling factor *α* = 1. The result is shown in Figure 12. This result infers, assuming *F* (*s*) = −1 that CORT dosage given at the time when the physical activity completed would have counteracted the temperature variations.

**Figure 12:**
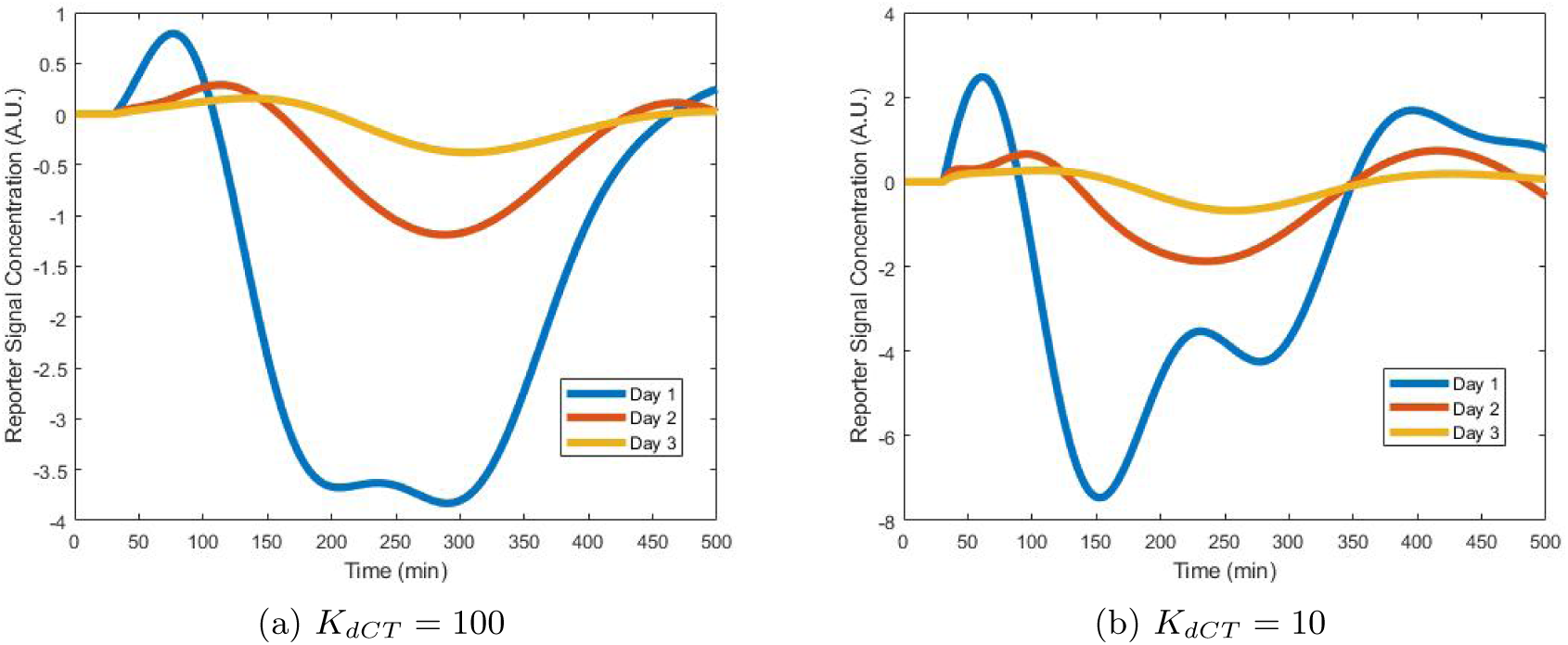
Through the use of the model the trajectory of cortisol contribution that is mapped to the temperature portion of the cardiovascular system status can be estimated. Two values of the PD control law are evaluated for different values of the derivative term of cortisol, with the other values of the controller kept constant at K_pT_ = 1, K_dT_ = 100, and K_pC_ = 1. It is seen that the shape of the concentration of cortisol follows along the lines of the observed temperature response but can be shifted in the later response stage by the derivative action. Although not shown, a similar effect is seen in the earlier stage of the response by altering K_dT_. A similar analysis can be made with the fatigue response, and the total result would be associated with the total cortisol fluctuation.

In Figure 12, the recurrent theme of the oscillatory nature of fever is seen in the responses in Day 1, relative to Day 2 and Day 3, when the vaccine had time to clear. The interference with the system dynamics, and the subsequent inefficiencies of the temperature control system are apparent. Thus, the information gathered in response to such excitation is useful in identifying the model equations of the temperature feedback system, which can subsequently be used for further analysis and predictive purposes.

To apply the model in behavior modification, one question that can be assessed using the model response is whether the observed temperature fluctuations of the case study due to the abrupt cessation of exercise would have been reduced had there been an extended cool down period in which the exercise was gradually reduced. This would seem to be the case since the frequency response of the closed-loop model indicates certain peaks which if excited would result in larger oscillatory responses, and therefore if the excitation can be kept to low frequencies, then less peak temperature fluctuation should be expected. This turns out to be the case, as seen in Figure 13, which compares the temperature fluctuation observed for Day 2, when the exercise was immediately stopped, in comparison to the temperature fluctuation had the exercise been gradually ramped down for the full period of observed nominal temperature oscillation. It is noted that temperature deviations during the exercise period were not observed because it was slow relative to the cessation of the exercise, and hence essentially a ramp, which can be modeled as 1*/s*^2^ that has significantly more frequency roll-off and attenuation then the step response of 1*/s*.

**Figure 13:**
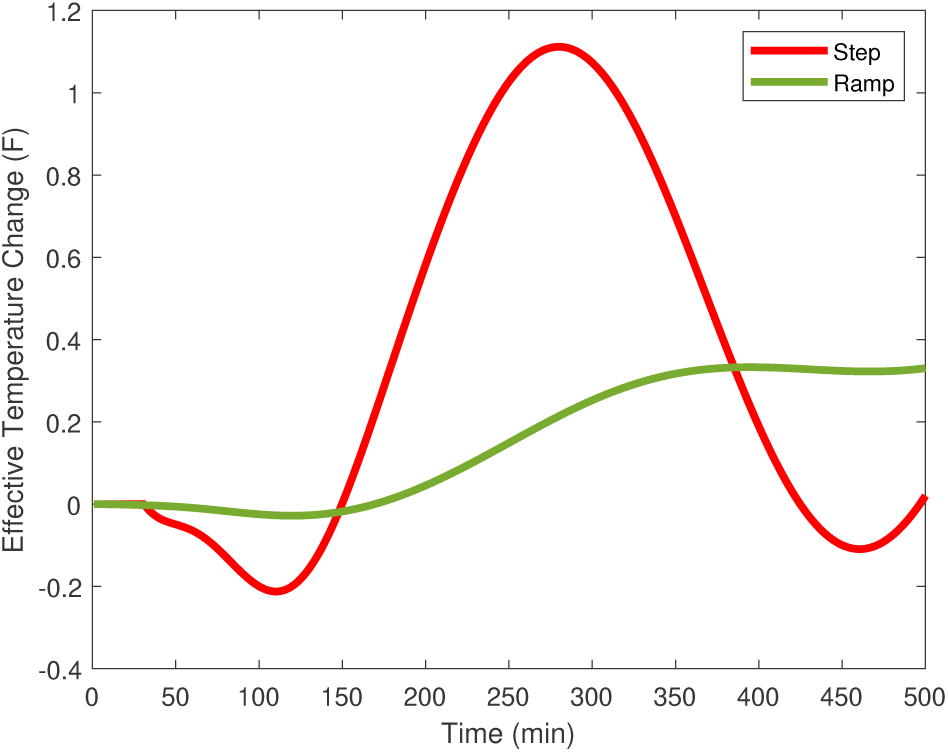
The model has predictive capability in optimizing and planning voluntary activity to compensate for sub-optimal situations. In this example, the temperature deviations are minimized if the cool down period after exercise were to have been conducted over an extended period, represented as a slow ramp down in activity, relative to just stopping the exercise regime, represented as step-down in the case study. The slow ramp down avoids the excitation of resonant frequencies in the dynamic response, which are present due to the vaccination altering the inherent dynamic response, pushing the system closer to instability in inducing oscillatory temperature modes.

## 3 Discussion

The molecular model comparing CORT and PGE2 in terms of the positional arrangement of the functional groups relative to GR is highly suggestive of competitive inhibition preventing normal systematic response of the hypothalamus. There is a near match of CORT and PGE2 in hydrogen bonding configuration at the LBD of GR, which creates an internal hydrophobic core, thereby achieving stability for each configuration and perhaps slightly favoring PGE2 due to the enhanced hydrogen bonding at one functional group. However, the slight discrepancy between the two is likely responsible for CORT being expressive of its DNA target gene, while that associated with PGE2 is non-expressive, hence dysfunctional.

The mathematical model suggests that the autonomic response is a function of not only the input neuroreceptor signal, but also is a function of the CORT concentration, and thus the response is co-dependent upon both neural and neuroendocrine signals. An immediate response is possible from the neural signal but is modulated over time by the CORT concentration, creating a fast response in conjunction with slow response. However, if the CORT signal is disrupted due to competitive inhibition at the GR for example, the overall effectiveness of the closed-loop response is greatly impacted. Essentially, the hypothalamus is presented with a faulty and incorrect input signal for which the system was designed - by analogy, this is a common problem in robust process control design for industrial system controllers or aeronautical controllers, in which the incoming measurement is not representative of the true system performance. In this case, CORT concentration is actually higher than that which is measured by the hypothalamus due to competitive inhibition of the GR with PGE2.

This misrepresentation of CORT from inhibition at the GR thereby induces dysfunction of the hypothalamus resulting in a range of symptoms seen in illnesses, such as body temperature fluctuations, fatigue, respiratory problems, and many other faulty autonomic responses. Thus, one signal error is responsible for many symptoms, and the different sequences and levels of response are dependent upon the nature of the PGE2 levels, as well as the dynamics of the load function and sensitivity of the gain functions of the hypothalamus. This can be counter-acted by reducing the levels of the conflicting chemical, or by diluting its effect by the addition of CORT, or by reducing the load inputs which will change the autonomic function.

The case study involving temperature fluctuation and fatigue also provides insight into fever, which is generally a most feared symptom. It indicates that fever is rather a natural response due to misprocessing at the hypothalamus because of an inaccurate application of CORT levels due to competitive inhibition at GR. That is, the neural and neuroendocrine response are not in synchronization. The temperature control system appears to be a highly tuned system capable of quick resolution of slow variations in exogenous inputs, however it shows a resonant frequency and thus is not robust to mismatch between nominal and infected system performance.

The interaction of the control loops is also quite interesting as shown in the derivation of the mathematical model, which naturally shows how the nervous, cardiovascular, immune systems are linked, and connected through the endocrine system. Moreover, other body systems such as the respiratory system are coordinated within the framework through the general parameterization of *X*(*s*). It indicates that the body systems are representative of an overall controller that is a highly integrated multi-input, multi-output (MIMO) system at the neural level coordinated with a single-input, multi-output (SIMO) system at the neuroendocrine level.

While it was necessary to study dysfunction, the work provided a way of understanding the functionality of the hypothalamus during normal operating conditions. Cortisol is essentially an indicator of the status of the cardiovascular system which is an effective linker of all body systems, and thus it is not surprising that the hypothalamus in making autonomic decisions on how best to control the internal systems, would use a measurement of the cardiovascular system in quantifying the response. It highlights the importance of maintaining a healthy cardiovascular system such that all other body systems can achieve optimal performance.

The work is useful for evaluating body system performance in the presence of infectious disease, developing diagnosis, and can be used for predictive purposes in developing new vaccines. It can be used to optimize behavior and drug treatment plans to minimize the symptoms of illnesses.

